# Functional characterization of *Ralstonia insidiosa*, a bona fide resident at the maternal-fetal interface

**DOI:** 10.1101/721977

**Authors:** Lindsay A. Parnell, Graham G. Willsey, Chetanchandra S. Joshi, Yin Yin, Matthew J. Wargo, Indira U. Mysorekar

**Author notes:** To whom correspondence should be addressed: Indira U. Mysorekar, Ph.D., Washington University School of Medicine, Depts. of Obstetrics and Gynecology; Pathology and Immunology, 660 S. Euclid Ave., St. Louis, MO 63110.

## Abstract

Controversy about whether there are microbes in the placenta and if they have any functional importance during pregnancy and for neonatal health is ongoing. Previous work has demonstrated that the basal plate (BP), comprising maternal and fetal derived cells harbors intracellular bacteria. 16S sequencing and bacterial species-specific analysis of term placentas revealed that the gram-negative bacillus *Ralstonia insidiosa*, native to aqueous environments and an effective biofilm promoter, comprises the most abundant species in the BP. Here, we demonstrate whether *R. insidiosa* cells home to a particular niche in the BP, how they may arrive there, and whether they are associated with adverse outcomes. We developed methods to detect and study cell-specific localization of *R. insidiosa* using ex vivo and in vitro models. Additionally, we studied potential routes of *R. insidiosa* entry into the placenta. We show that *R. insidiosa* is a bona fide resident in human placental BP. It can access trophoblast cells in culture and within basal plate tissues where it localizes to intracellular single-membrane vacuoles and can replicate. However, the presence of *R. insidiosa* does not cause cell death and does not induce a pro-inflammatory immune response suggesting that it is not harmful in and of itself. Finally, we show that in a pregnant mouse model, *R. insidiosa* traffics to the placenta via the intrauterine route but does not induce preterm labor or preterm birth. Together, our findings provide a foundation for understanding non-pathogenic placental cell-microbe interactions and the functional importance of *R. insidiosa* in placental health and physiology.

## Introduction

Throughout pregnancy, the fetus relies on the placenta for communication with the mother and transport of nutrients and wastes. Additionally, the placenta provides a barrier against ascending and hematogenously transmitted infections. Most microorganisms cannot penetrate the placenta, though some can bypass placental defenses, leading to local or systemic immune activation and adverse outcomes such as preterm birth or fetal death ^**1**^. Thus, the healthy uterus has long been considered a sterile environment ^**2**^.

Contrary to this paradigm, bacterial DNA has been identified in the amniotic fluid and meconium, suggesting that the womb may not be sterile after all ^3–17^. More recently, studies have detected bacteria in term placentas ^**11,18–24**^. For example, our histological analysis revealed the presence of both gram-positive and gram-negative intracellular bacteria, sometimes in biofilm-like clusters of bacilli, in the human placental basal plate (BP), which contains both fetal and maternal cells ^23^. Seferovich and colleagues further validated the presence of bacteria from term and preterm placentas, without evidence of maternal infection, using 16S in situ hybridization, histologic staining methods, and clinical culturing methods ^22^. Additionally, several investigators have used culture-independent sequencing methods to identify bacterial DNA in term and preterm human placentas ^18,22,25,26^ and other fetal or maternal conditions including low birth-weight ^27^, preeclampsia ^28^, maternal obesity ^29^, and gestational diabetes ^30^.

We previously performed 16S ribosomal RNA (rRNA) sequencing to interrogate the microbial profiles at various sites in full term placentas from uncomplicated pregnancies. These sites include the maternal compartment, comprised of the BP, and the fetal compartment, comprised of the placental villi (PV) and fetal membranes (FM). We found that *Ralstonia insidiosa* (*R. insidiosa*), a gram-negative, aerobic, motile bacillus, is most prevalent in the BP ^19^. *R. insidiosa* was first isolated from the sputum of cystic fibrosis patients and can be isolated from immunocompromised patients in hospitals ^31,32^. However, this bacterium is also found in pond water, soil, sludge ^33^, fresh-cut produce facilities ^34^, water distribution systems ^35^, and dispensed water used for bottled water ^36^. Because *R. insidiosa* can form biofilms ^34,37^, is multi-drug resistant, and can survive sterilization methods ^38^, it may pose a health threat to some individuals ^39^.

Here, we sought to determine whether or not *R. insidiosa* is present at the maternal-fetal interface and its potential relevance in this niche. We report that we can identify *R. insidiosa* by fluorescent in situ hybridization (FISH) in human placental BP biopsies. Additionally, we show that *R. insidiosa* can enter and replicate within human trophoblasts ex vivo. Further, we show that *R. insidiosa* replicates in a model of extravillous trophoblasts (EVTs) in an in vitro setting, but it does not cause cell death or activate a pro-inflammatory immune responses. Finally, we show that *R. insidiosa* can enter the mouse placenta when delivered via the intrauterine route, though it does not cause preterm labor. Our findings support the idea that *R. insidiosa* can colonize the placenta but is likely not a threat to placental health. Here, we report *R. insidiosa* as a bona fide resident in the maternal-fetal interface.

## Results

### *R. insidiosa* is detected in term BP specimens

Due to lack of availability of antibodies to detect *R. insidiosa* in preserved tissue, we developed FISH probes that could be used on placentas. We used the DesignProbes package in Decipher ^40^, which assesses specificity and calculates potential cross-hybridization species, to design a species-specific probe for *R. insidiosa*. The specificity of this probe was confirmed by growing the *R. insidiosa* reference strain (MJ602) in culture and performing FISH with either a bacterial-domain probe (Bacteria – EUB338-CY3) and a species–specific *R. insidiosa* probe (**Figure S1 A-D**), a *R. insidiosa* probe alone (**Figure S1 E-H**), or a non-sense negative control probe (Non-bacterial - NON-EUB338) and bacterial probe (**Figure S1 I-L**). The species-specific probe, with and without the application of the bacterial probe, hybridized as expected using the reference *R. insidiosa* strain. Failure to detect the non-specific nonsense probe confirmed the functionality of the bacterial probe for our study (**Figure S1 I-L**). The probes and the bacteria used in this study are presented in **Tables S1 and S2**, respectively.

Next, we tested whether *R. insidiosa* could be detected in BP biopsies from our previous study (**Table 1**). 11 BP samples from term pregnancies were tested. We also stained the samples with an antibody to histocompatibility complex-G (HLA-G), a marker of EVTs within the BP (**Figure 1 A-B**). Bacteria were detected in the BP with both the bacterial probe (**Figure 1 C, H**) and the *R. insidiosa*-specific probe (**Figure 1 D, I**) in two cases. Further, composite images including differential interference contrast (DIC) imaging and DNA staining (**Figure 1 E-L**) revealed that the probes hybridized to bacterial DNA within large nucleated trophoblast cells. Together, FISH analysis confirmed the presence of bacterial species in the maternal BPs and delineated that *R. insidiosa* was one of the bacterial species in this region.

**Table 1.** Summary maternal donors of clinical BP specimens.

| Case no. | No. of *R. insidiosa* reads<sup>a</sup> | Gestational age at delivery (weeks & days) | Maternal age | Maternal race | Spontaneous labor | Indicated birth | Mode of delivery |
| --- | --- | --- | --- | --- | --- | --- | --- |
| 1 | 498 | 39 & 2 | 32 | White | No | Yes | C-section |
| 7 | 3457 | 40 & 2 | 22 | Black | No | Yes | Vaginal |
| 8 | 465 | 39 & 5 | 28 | Black | Yes | No | C-section |
| 9 | 2238 | 39 & 2 | 21 | Black | Yes | No | Vaginal |
| 13 | 2469 | 37 & 3 | 30 | Caucasian | No | Yes | Vaginal |
| 14 | 6418 | 40 & 0 | 35 | Asian/PI | Yes | No | Vaginal |
| 15 | 2125 | 40 & 4 | 26 | Black | Yes | No | C-section |
| 16 | 1482 | 37 & 6 | 19 | White | Yes | No | Vaginal |
| 17 | 1458 | 39 & 6 | 30 | White | No | Yes | Vaginal |
| 19 | 0 | 37 & 1 | 30 | Caucasian | No | Yes | C-Section |
| 20 | 0 | 40 & 3 | 32 | White | Yes | No | Vaginal |

<sup>a</sup>Sequencing reads for cases were obtained from <sup>19</sup>.

**Figure 1.**
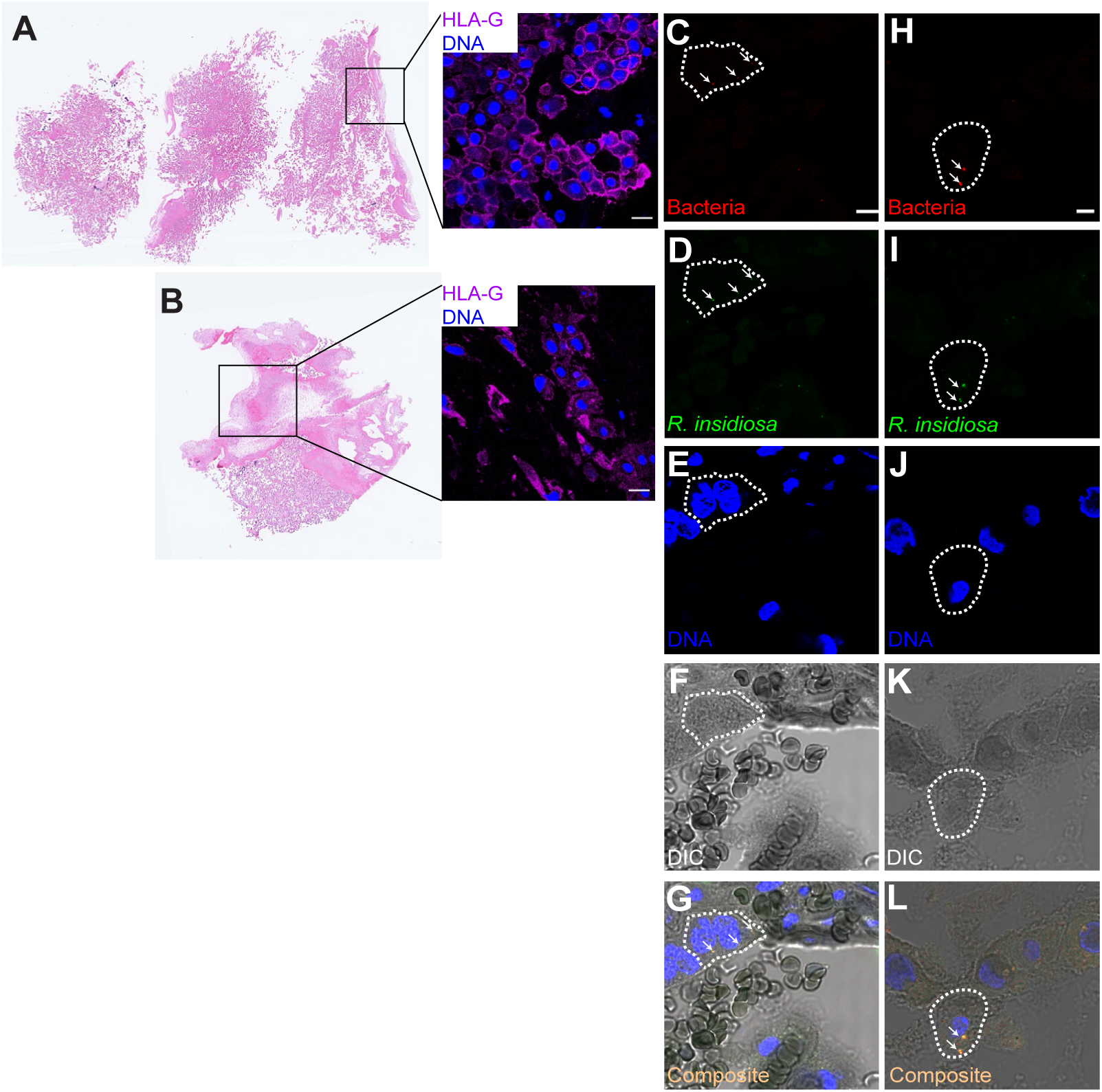
*R. insidiosa* is detected in clinical BP samples positive for *R. insidiosa* by 16S sequencing. (A-B) Hematoxylin and eosin staining of 2 clinical term BP specimens from cases positive for *R. insidiosa*. Specimens demonstrate the presence of the EVT-specific marker HLA-G (purple) (Magnification 40x oil, scale bars: 20 µm). (C-G) FISH performed on a BP specimen from the case presented in (A), simultaneously hybridized with the (C) bacterial probe (red), (D) the R. insidiosa-specific probe (green), (E) DAPI (DNA-blue). (F) Digital interference (DIC) and (G) composite images are also displayed. White arrows indicate areas of colocalized bacteria and R. insidiosa probes on the periphery of host nuclei. (H-L) Similar observations are shown in a BP specimen from the case presented in (B) (Magnification 63x oil, scale bars: 5 µm).

### *R. insidiosa* can colonize BP explants and EVTs

We previously reported that, when added to BP explants cultured ex vivo, gram-positive and negative bacteria replicate and aggregate within EVTs ^41^. These epithelial cells extravasate from the placental villi into the BP to anchor the fetal compartment to the maternal side of the placenta. This event both remodels spiral arteries and facilitates blood circulation within the intervillous space ^42^. We wondered whether *R. insidiosa* could also invade and replicate within BPs, and specifically, within EVTs. In addition to the WT *R. insidiosa strain* (MJ602), we generated yfp-expressing *R. insidiosa* strain (GGW102) to localize *R. insidiosa* within BP explants (**Table S2**). Species-specific FISH probes effectively hybridized to GGW102 (**Figure S2 A-L**). Additionally, we confirmed that it expressed yfp using a panel of YFP-specific qPCR primers generated commercially (**Figure S2 M; Table S3**). *R. insidiosa* WT (MJ602) and yfp-tagged (GGW102) isolates were cultured for over the course of 24 hours to monitor growth patterns (**Figure S3**). We did not see a significant difference in the log colony forming units (CFUs) between isolates after 4 hours, indicating that MJ602 or GGW102 could be used interchangeably to assess *R. insidiosa* growth.

Fresh biopsies from placentas from term uncomplicated pregnancies delivered vaginally or by Caesarean section were incubated with 1×10^8^ CFUs of *R. insidiosa* yfp-strain (GGW102) for 8 hours. The explants remained morphologically intact, suggesting that *R. insidiosa* caused no major cell damage (**Figure S2 A-B).** Species-specific FISH revealed *R. insidiosa* within cells adjacent to large nuclei in the BP explant (**Figure 2 C**). Using transmission electron microscopy targeting *R. insidiosa* cells (**Figure 2 D**) and *R. insidiosa*-challenged explants (**Figure 2 E-F**), we noted that single BP cells often contained multiple bacteria within single-membrane vacuolar compartments.

**Figure 2.**
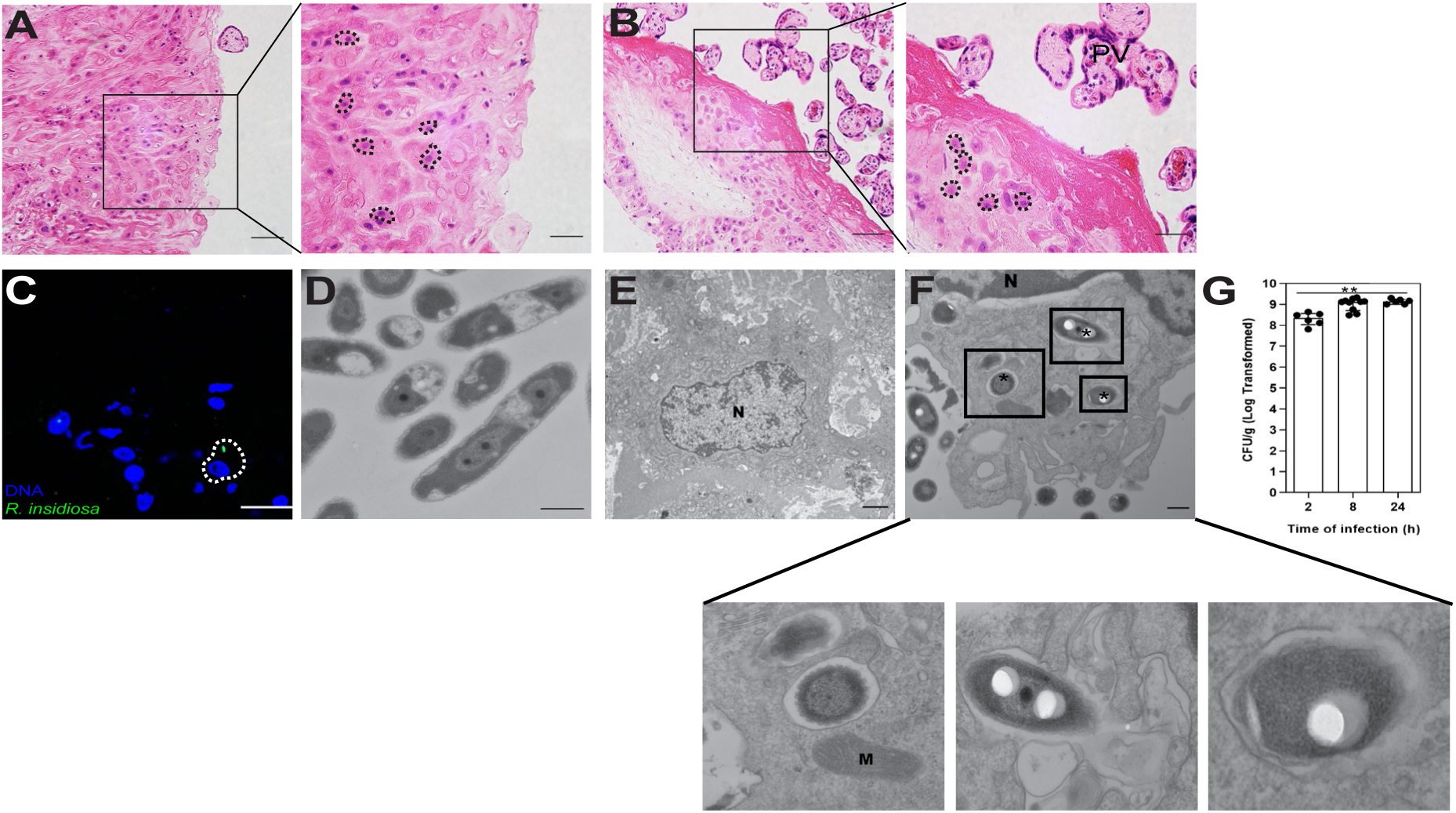
*R. insidiosa* colonize basal plate explants. Hematoxylin and eosin staining shows the histological landscape of (A) *R-insidiosa*-challenged and (B) unchallenged BP explants (8 h) (magnification 20x, scale bar: 100 µm). Magnified regions show intact mononuclear extravillous trophoblasts (dashed lines) (magnification 40x, scale bars: 100 µm). (C) Explants hybridized with *R. insidiosa* probe (*R. insidiosa*-green) and DAPI (DNA-blue). *R. insidiosa* exhibits a rod-shaped morphology (magnification 63X oil, scale bars: 20 µm). Representative TEM images of (D) pure bacterial culture of *R. insidiosa* (MJ602) at 16 h (magnification 15000x, scale bar: 0.5µm) and (E) a single cell within an unchallenged BP explant (magnification 2500x, scale bar: 2 µm). (F) BP explants challenged with *R. insidiosa* at 2 h (magnification 10000x, scale bar 0.5 µm). Enlargements of box regions in (F) indicate R. insidiosa (*) within single-membrane vacuolar compartments. (G) *R. insidiosa* grows on or within BP cells over time as indicated by CFU analysis (n = 3, median: 2 h = 8.36 Log cfu/g, 8 h = 9.13 Log cfu/g, 24 h = 9.14 Log cfu/g; Kruskal-Wallis statistic= 11.3, p = .0010). Graphs present the median and interquartile range. P-values were calculated using Kruskal-Wallis with Dunn’s Multiple Comparison Test (*, p < 0.05 and **, p < 0.005).

Next, we wondered whether *R. insidiosa* proliferates within the BP cells. Because *R. insidiosa* is resistant to gentamicin (data not shown and^38^), the standard gentamicin protection assay could not be used to assess intracellular bacterial replication. Instead, we incubated the explants with 1×10^8^ CFU of *R. insidiosa* for 2, 8, and 24 hours. At each time point, explants were extensively washed, homogenized, and quantified for adhered and invaded bacteria using CFU analysis. We found that the *R. insidiosa* CFUs increased approximately ten-fold from 2 to 24 hours, suggesting that the bacteria replicated either within or on BP explant cells (**Figure 2 G**). In conjunction with TEM, this indicates that *R. insidiosa* replicates intracellularly within BP cells.

To directly address whether *R. insidiosa* could invade EVTs, we used the JEG-3 choriocarcinoma line. JEG-3 cells have a similar gene expression profile to normal EVTs ^43,44^ and have been used as a model to study pathogen-EVT interactions in the placenta ^45^. We challenged JEG-3 cells with *R. insidiosa* at multiple doses and quantified bacterial invasion and adhesion. Although *R. insidiosa* could adhere to and invade JEG-3 cells, it did not do so in a dose-dependent manner (**Figure 3 A**). Similar to *R. insidiosa*-challenged BP cells, TEM analysis of JEG-3 cells revealed single *R. insidiosa* within single-membrane vacuoles (**Figure 3 B-F**). In some cases, multiple *R. insidiosa* rods existed within a single vacuole. These vacuoles were reminiscent of Listeria monocytogenes-containing vacuoles recently reported by Kortebi and colleagues ^46^ (**Figure 3 G-I**).

**Figure 3.**
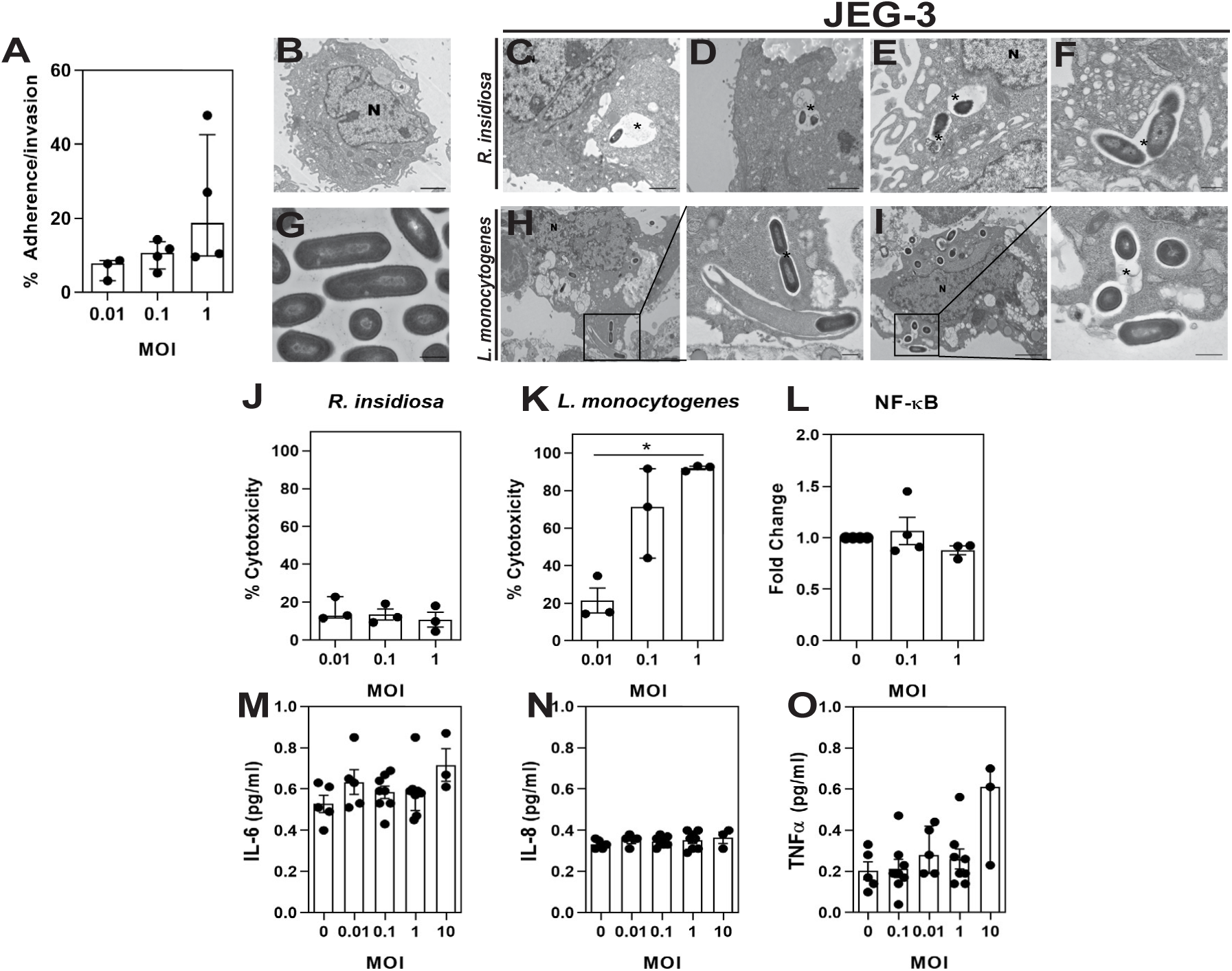
*R. insidiosa* colonize JEG-3 cells and does not elicit pro-inflammatory responses. (A) Percent adherence and invasion of R. insidiosa at MOIs of 0.01, 0.1, and 1 (Kruskal-Wallis statistic: 4.89, p = 0.0836). Representative TEM images of (B) unchallenged JEG-3 cells and (C-F) *R. insidiosa*-challenged cells (n=3) at 8h. The * indicates *R. insidiosa* rod-shaped morphologies existing within single-membrane vacuolar compartments (magnification: A-2500x, B-5000x, C-6000x, D-7500x, E-12000x). (C, E) Bacteria existed in these compartments singly and, at times, (D, F) two were present in a single compartment. Representative TEM images of pure bacterial culture of *L. monocytogenes* (EGD) at 16 h (magnification 20000x, scale bar: 0.5µm). (H, I) Similar to *R. insidiosa, L. monocytogenes* may exist intracellularly within single-membrane vacuolar compartments as indicated by the * symbol (magnification: H-3000x, I-5000x). Boxed regions are magnified regions of and (I) (magnification: 12000x; 20000x). (J) Percent cytotoxicity of JEG-3 cells following *R. insidiosa* exposure for 8 h indicate lack of cell death at various MOIs (Kruskal-Wallis statistic = 1.42, p = 0.543) compared to (K) *L. monocytogenes* (Kruskal-Wallis statistic =6.49, p = 0.0107). Challenge with *R. insidiosa* at various MOIs (8 h) indicates a lack of expression of pro-inflammatory transcription factor (L) NFκ-B (Kruskal-Wallis statistic = 2.98, p =0.233) and lack of secretion of pro-inflammatory cytokines including (M) IL-6 (Kruskal-Wallis statistic = 5.74, p =0.219), (N) IL-8 (Kruskal-Wallis statistic =2.63, p = 0.622), and (O) TNF-α (Kruskal-Wallis statistic = 6.19, p =0.185). P-values were calculated using Kruskal-Wallis with Dunn’s Multiple Comparison Test (*, P <0.05). TEM images scale bars: (A-C, G-H) 2 µm, (D-F) 0.5 µm.

To determine the impact of *R. insidiosa* on EVTs, we first assessed cell viability by quantifying lactate dehydrogenase, which is released upon cell death in media. Whereas *R. insidiosa* caused no JEG-3 cell death (**Figure 3 J**), challenge with *L. monocytogenes* did cause cell death in a dose-dependent manner (**Figure 3 K**). Next, we asked whether EVTs mount inflammatory responses when exposed to *R. insidiosa*. To answer this question, we challenged JEG-3 cells with *R. insidiosa* for 8 hours and then evaluated expression of the pro-inflammatory immune marker NF-κB and the secretion of the pro-inflammatory cytokines IL-6, TNF-α, and IL-8, which are upregulated upon activation of the NFκB pathway ^47^. None of these were upregulated in *R. insidiosa*-challenged JEG-3 cells (**Figure 3 L-O**). Together, we conclude that *R. insidiosa* can exist within EVTs without inducing death or pro-inflammatory responses.

### *R. insidiosa* is detectable and cultivable in the murine placenta

How bacteria reach the placenta has not been fully determined. Several studies have reported that non-pathogenic microbes in the placenta are most similar to those found in the oral mucosa ^18,29,48^. For example, the oral bacterium *Fusobacterium nucleatum* has been isolated from the placenta and amniotic fluid ^49^. Thus, we investigated possible sources of *R. insidiosa* within the placenta. We performed meta-analysis of the Human Microbiome dataset and found *Ralstonia spp.* in niches associated with the oral mucosa including the hard palate, saliva, and throat (**Figure 4 A**). *Ralstonia* was also detected in the anterior nares and left and right retroauricular crease and antecubital fossa (data not shown). In contrast, *Ralstonia* was found at low abundance in stool.

**Figure 4.**
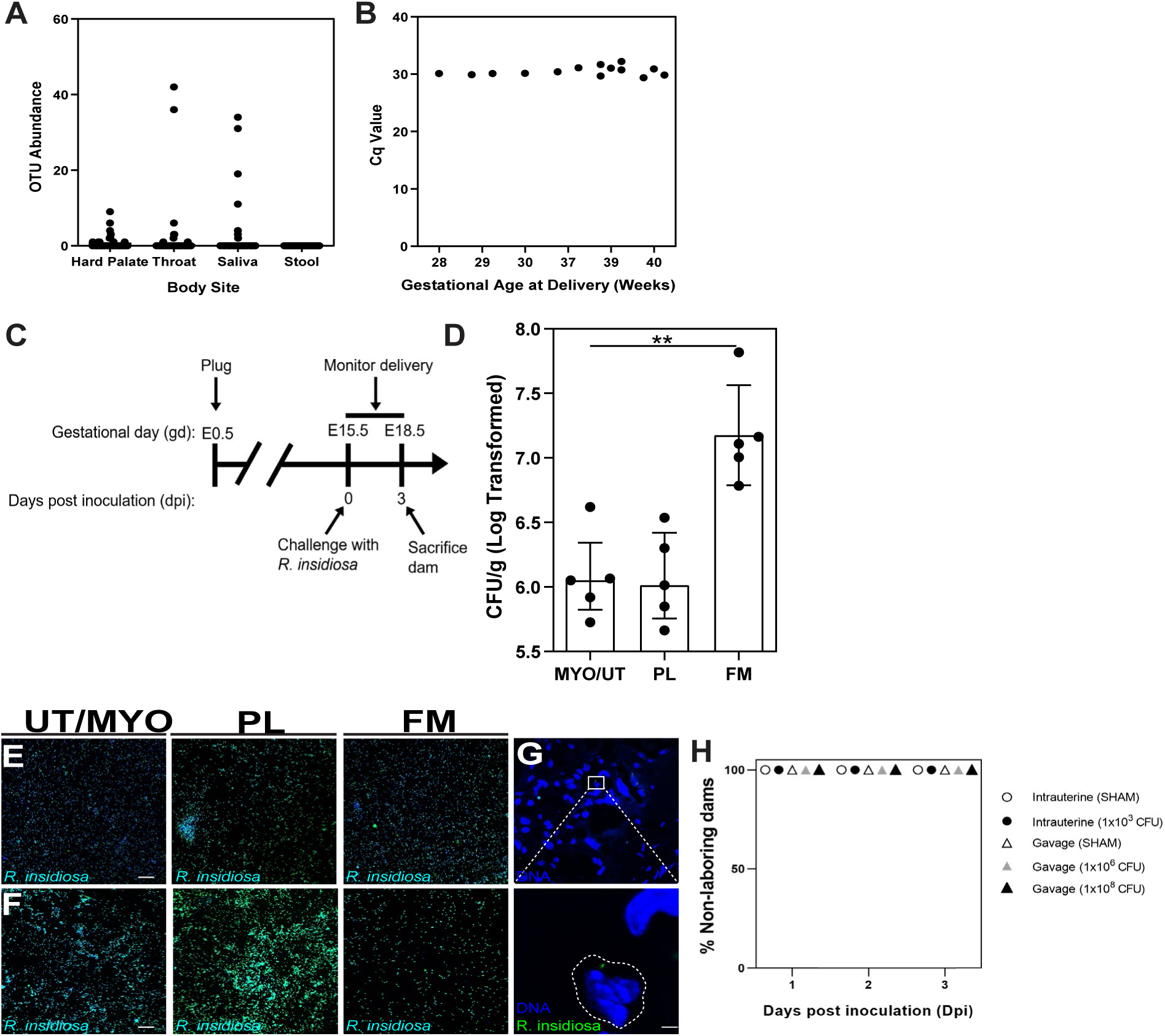
*R. insidiosa* colonize the murine placenta following intrauterine injection. (A) Meta-analysis of HMP dataset demonstrating the abundance of *Ralstonia*-specific operational taxonomic units (OTUs) in various niches. (B) *R. insidiosa*-specific qpcr of BP biopsies indicating the detection of *R. insidiosa* as early as 28 weeks. (C) Overview of *R. insidiosa* challenge in WT dams by oral gavage and intrauterine injection. (D) Mean-bacterial load (Log transformed cfu/g) in the placenta, uterus/myometrium (UT/MYO), and fetal membranes (FM) per dam (n=5 median: UT/MYO = 6.052 log cfu/g, PL= 6.014 log cfu/g, FM= 7.109 log cfu/g, Kruskal-Wallis Statistic=9.429, P = 0.0024) challenged by intrauterine injection. P-values were calculated using Kruskal-Wallis with Dunn’s Multiple Comparison Test (*, P < 0.05; **, P < 0.005). (E-F) FISH was performed on a single colony cultivated from the UT/MYO, PL, and FM of a representative fetal units from 2 separately challenged dams. The representative confocal images show the cultivated *R. insidiosa* (green) and bacterial DNA (blue). (G) Representation of mouse placentas hybridized with the *R. insidiosa* probe (green) and DAPI (DNA-blue), along with a zoomed image of the boxed region (magnification 63x oil, scale bar: 2µm). FISH images were captured using the Carl Zeiss Confocal Microscope. (H) % of mice without signs of labor post-inoculation until sacrifice.

Given that *Ralstonia spp.* have been detected in oral niches and has been found in fresh-cut produce facilities ^34^, water distribution systems ^35^, and bottled water ^50,51^, we hypothesized that this bacterium could reach the placenta via the oral route. To test this idea, we administered 1×10^6^ or 1×10^8^ CFU of *R. insidiosa* to pregnant wild-type (C57BL/6) mice at gestational day 15.5. This strain of mice delivers at 19 days ^52^, so this would be roughly equivalent to 32 weeks’ gestation in human pregnancy, a time when we detected *R. insidiosa* in human BP biopsies (**Figure 4 B**). We sacrificed the dams at gestational day 18.5 (**Figure 4 C**). However, we were unable to detect measurable *R. insidiosa* CFUs in the uteri/myometrium (UT/MYO), fetal membranes (FM), or placentas (PL). We observed no significant differences in placental weight, fetal weight, fetal width, or fetal length between control mice and those that were administered *R. insidiosa* (**Figure S4 A**). Further, none of the mice that received bacteria delivered early (**Figure 4 H**). Thus, *R. insidiosa* administered orally did not adversely affect delivery.

Intrauterine bacterial colonization is associated with preterm birth ^53^. Thus, we asked whether *R. insidiosa* delivered directly to the uterus would colonize the placenta and possibly lead to preterm labor. At day 15.5, pregnant dams were administered two intrauterine injections totaling 1×10^3^ CFUs and were monitored for signs of labor until day 18.5. Dams were sacrificed at 18.5 and the UT/MYO, FM, and PL were collected and assessed for CFUs (**Figure 4 C**). We were able to recover 1×10^6^ to 1×10^7^ *R. insidiosa* from all of these tissues (**Figure 4 D**), indicating that *R. insidiosa* grew over 72 hours. *R. insidiosa* was most abundant in the FM. Additionally, we detected *R. insidiosa* in all of these tissues by *R. insidiosa*-specific FISH analysis (**Figure 4 E-G**). Despite the robust *R. insidiosa* growth in the myometrium, placenta, and fetal membranes, *R. insidiosa* did not stimulate preterm labor (**Figure 4 H**) or affect placental weight, fetal weight, fetal width, or fetal length (**Figure S4 B**). Together, these findings suggest that *R. insidiosa* may reach the placenta via the intrauterine route but that it does not cause any adverse pregnancy outcomes on its own.

## Discussion

Whether there are microbiota (live or simply DNA signatures) in the maternal-fetal interface remains mired in controversy. In the recent years, there has been a large number of studies, providing both sequencing and histological evidence, indicating microbial signatures ^3,11,18,19,22,23,29^, while other studies have reported these signatures as likely DNA contaminants originating from the sample collection and/or the DNA extraction process, rather than the placenta itself ^54–57^. Placenta location remains a key variable in the aforementioned studies given that that BP, PV and FM have distinct physiology and function. However, many studies interrogating the placental microbiota have not focused on the BP itself, home to fetal EVTs and immune cells. Rather, many studies have focused on either the PV alone, which is in direct contact with maternal blood and the site of nutritional transport and gas exchange, or do not distinguish between the BP and PV when sampling.

We found a high prevalence of *R. insidiosa*, a waterborne environmental bacterium, in human BPs using multi-variable region 16S sequencing approach ^19^. Other investigators have also identified *Ralstonia spp*. in the placenta ^25,58^ and amniotic fluid ^3^. However, some studies suggest *Ralstonia. spp*. detected by sequencing represent contaminants originating from DNA extraction kits or other sources ^54–56,58,59^. In this report, we identify *R. insidiosa* as a bona fide resident at the maternal-fetal interface by using species-specific FISH probes to confirm and localize *R. insidiosa* within BP tissue specimens at term.

A recent study from Theis and colleagues provide guidelines for how to define placental microbiota ^58^. *First*, specimens must demonstrate a distinct microbial signal compared to negative controls in order to exclude contamination as the source of bacterial DNA. In our previous study, we employed a series of pre- and post-16S sequencing steps to minimize risk of analyzing false microbial signals ^19^. Using 16S qPCR, for example, we detected microbial signals above background and only analyzed the microbial composition of those samples. Using species-specific qPCR we also demonstrated the presence of *R. insidiosa* DNA in the BP and verified its absence in negative controls ^19^.

*Second*, the guidelines recommend that the detected microbial signals must be validated using multiple methods ^58^. In the current study, we present the first use of *R. insidiosa*-specific probes to localize the bacteria intracellularly within term human BP specimens, which we previously established were positive for *R. insidiosa* by 16S sequencing and species-specific qPCR ^19^. These probes predominately bind to 16S rRNA and the bright staining suggests a large intact rRNA pool, which is typically a hallmark of viability. *Finally*, the guidelines suggest that it must be “ecologically plausible” for the detected microbes to reside in the placenta ^3,58^. In this report, we show that *R. insidiosa* replicates within human placental samples, human BP explants cultured ex vivo, and EVT-like JEG-3 cells in culture. Further, intracellular *R. insidiosa* bacteria were found in single-membrane bound compartments that closely resembled Spacious Listeria-containing vacuoles (SLAPs) ^60^ or Listeria-containing vacuoles (LisCVs) ^46^. Using JEG-3 cells, Kortebi et al., showed that *L. monocytogenes* persist in LisCVs, which are a unique, acidic vacuolar compartments that are not degradative ^46^. In JEG-3 cells, we observed *R. insidiosa* within clusters or individually within each vacuolar compartment. We speculate that *R. insidiosa* may also utilize similar non-degradative compartments to adopt a “quiescent lifestyle” ^46^ within EVTs. Consistent with this, we found *R. insidiosa* is not harmful to EVTs as it does not induce responses associated with pathogenicity — including cell death or heightened pro-inflammatory responses.

How *R. insidiosa* may reside in EVTs remains elusive. We previously showed that uropathogenic *E. coli* (UPEC) persists in bladder epithelial cells by co-opting the feritinophagy pathway and utilizing bioavailable iron for intracellular growth ^61^. Similar to UPEC, some *Ralstonia spp.*, including specific strains of *R. picketti* ^62^ and *R. eutropha* ^63^, either demonstrate siderophore activity or harbor gene clusters with putative siderophore functions, suggesting that they have the capacity to chelate iron for growth. However, it is not clear whether *R. insidiosa* triggers autophagy or scavenges intracellular iron in order to persist in EVTs. Further studies are needed to determine the molecular mechanisms by which R. insidiosa resides in EVTs.

*Ralstonia spp. are* native to aqueous environments and have been isolated from variety water sources consumed by humans ^35^. Using human microbiome database, we showed that *Ralstonia spp.* are detectable in a variety of oral niches. However, we were not able to cultivate or detect *R. insidiosa* from placentas by this route. Notably, mouse studies that have traced oral bacteria to the placenta have used intravenous injections (such as tail vein injections) as a method of bacterial administration to model the bloodstream as a source of intrauterine and placental colonization ^48,49^. It is possible that *R. insidiosa* need to enter maternal circulation to access the placenta. Our results suggest that *R. insidiosa* did not enter the bloodstream when administered directly to the stomach via oral gavage. However, it is possible that *R. insidiosa was able to* traffic to the placenta via the oral route in extremely low amounts, but was undetectable using our cultivation methods.

Our studies provide the first in vivo evidence that viable *R. insidiosa* colonize and replicate within the murine placenta following a local administration via intrauterine injection. LPS alone administered by intrauterine injection at the similar gestational ages (14.5 to 15.5) has been shown to induce preterm labor (delivery at < 19 days) ^64^. We show that *R. insidiosa*, a gram-negative bacterium harboring LPS, does not induce preterm labor. Together, this demonstrates that presence *R. insidiosa* at the later stages of pregnancy is not associated with adverse pregnancy outcomes.

Using pregnant ewes, Zarate and colleagues recently demonstrated that maternal and/or fetal hypoxic stress later in pregnancy upregulates neuro-immune pathways associated with LPS stimulation ^65^. Under these conditions, this group detected live antibiotic-resistant *Staphylococcus simulans* in the placenta and fetal brain, suggesting that specific physiological changes associated with hypoxic stress, such as increased blood flow and barrier permeability, can promote the passage of maternal bacteria across the placenta and fetal blood-brain barrier ^65^. Similarly, maternal stress levels or other unknown physiological changes during pregnancy may explain the presence of *R. insidiosa* in the human placenta. As suggested by Zarate and colleagues, maternal antibiotic administration may also contribute to the transit of antibiotic resistant bacteria to the placenta. Thus, future studies must address how both external factors and maternal physiology lead to the presence of *R. insidiosa* in this niche.

In sum, our data indicate that it is ecologically plausible for *R. insidiosa* to be a bona fide placental microbial resident because it exists intracellularly within BP without causing adverse responses (ie. inflammation and cell death) and may translocate from the uterus to the placenta without inducing preterm birth.

## Materials and methods

### Study approval

This study was approved by the Washington University School of Medicine human studies review board (IRB ID 201012734) as described in ^19,41,66^. Informed consent was retrieved from all subjects used in this study. All animal procedures were reviewed and approved by the animal studies committee of the Washington University School of Medicine.

### Species-specific oligonucleotide probe design and specificity evaluation

A species-specific probe was designed to visualize *R. insidiosa* using the DesignProbes package in Decipher ^40^, which assesses specificity and calculates likely cross-hybridization of other species sequences. A universal bacterial probe (EUB338), complementary to conserved regions of the bacterial 16S rRNA gene was used as a positive control ^67^ and a nonsense probe (NON-EUB338) ^68^ was used as a negative control. All fluorescently labeled probes were synthesized commercially (Integrated DNA Technologies, Coralville) with the 5’ end labeled with either Cyanine-3 (CY3) or fluorescein (FITC). The probe specificity for MJ602 and GGW102 were evaluated in this study by whole-cell fluorescent in situ hybridization. Briefly, cultured bacteria were fixed with 4% paraformaldehyde for 1-3 hrs. Cells were washed with 50%, 85%, and 95% ethanol for 5 minutes and air-dried. Samples were pre-hybridized at 37 °C with hybridization buffer (0.6 M NaCl, 30% formamide. 10 mM tris-HCl, 0.25% SDS, 100 ug/ml salmon sperm, 1 mM EDTA, 1X Denhardt’s Solution) for 30 minutes. Samples were hybridized with EUB338-CY3 and *R. insidiosa*-FITC, EUB338-CY3 and Non-EUB338-FITC or *R. insidiosa*-FITC alone. Hybridized pellets were washed in 0.1X saline sodium citrate (SSC) and counterstained with DAPI (4’, 6’-diamidino-2-phenylindole) using ProLong Diamond Antifade Mountant with DAPI (Invitrogen, Eugene) to detect bacterial DNA.

### Placental BP specimens

Our previous study showed that 17 term cases had greater than 300 reads for *R. insidiosa* ^19^ and 11 of these cases were used for this study (Median = 2125 reads, IQR =1011 reads). BP biopsies from these cases were formalin-fixed at the time of delivery and paraffin embedded. A total 42 tissue sections were hybridized with *R. insidiosa*-FITC and EUB338-CY3 probes, 7 of which (n = 2 cases) were used as negative controls and were derived from cases with no detectable *R. insidiosa* by 16S sequencing. Sections were occasionally tested in parallel with the non-specific NON-EUB338 probe to ensure probe specificity. Subjective observation based on signal intensity and the overlap of the species-specific and eubacterial probe was used to detect the *R. insidiosa* signal. Additionally, sections underwent H&E staining as described in ^23^.

### Bacterial strains and growth kinetics

*R. insidiosa* strain MJ602 was isolated from the International Space Station potable water reclamation system and is stocked at the Johnson Space Center as *R. insidiosa* 130770013-1 ^69^. The YFP-expressing *R. insidiosa* strain GGW102 was generated by *attTn7* integration into MJ602 using co-electroporation of pUC18T-mini-Tn7T-Tp-eyfp (DQ493883) and pTNS2 (AY884833) followed by selection on LB agar with 100 g/ml trimethoprim using the strategy described in ^70^. Integrants were verified by PCR and positive YFP fluorescence. Reasoner’s 2 Agar (R2A) (EMD Millipore) and Reasoner’s 2 Broth (R2B) (EMD Millipore) were used for routine bacterial growth of *R. insidiosa. R. insidiosa* from the glycerol stocks were streaked onto R2A using a sterile loop and grown at 32 °C until small white round colonies formed. Single colonies were grown in R2B at 32 °C for the appropriate incubation periods. For growth curves of each strain, the optical density at 600 nm (OD_600_) was recorded every 4 hours over a 24-hour incubation period (n=6 colonies). Colonies grown on R2A and incubated at 32 °C overnight was used to enumerate colony-forming units (CFUs). To calculate CFUs, the following equation was employed: number of colonies x dilution factor/volume on the culture plate. Finally, CFUs were log-transformed and plotted against time or OD_600_ for each strain. *L. monocytogenes* was cultured as described in ^41^.

### BP explant culture

Placentas (n= 5) from term pregnancies delivered by vaginal or Caesarean section were collected within 1-4 hours of delivery and cultured as described previously ^41^ with the following modifications: BP explants were seeded on to either collagen I-coated 12-well plates and cultured in 1 ml of culture medium (DMEM with 10% fetal bovine serum) or 24-well plates and cultured in 0.5 ml of culture medium. Explants were challenged with 1×10^8^ colony forming units of *R. insidiosa* (GGW102) and centrifuged at 1500 RPM for 5 minutes followed by incubation for 8 hours. To quantify adhered and/or invaded bacteria, media was removed from each well and washed with sterile PBS. Explants from each well and time point were weighed and homogenized in 400 µL of PBS and plated on R2A. The log-transformed median CFU/g per well was calculated and plotted against time. Alternatively, explants were fixed in 4% paraformaldehyde (Fisher Scientific, Fair Lawn NJ) at 37 °C for 30 minutes to 1 hour and embedded in 2% agar (Sigma, St. Louis MO). Explants were processed in 3-5 micron sections. Sections underwent H&E staining, immunofluorescence, and fluorescent in situ hybridization.

### Fluorescent in situ hybridization

FPPE sections of placental BP biopsies and cultured explants were deparaffinized and hydrated. Sections were microwaved in 10 mM sodium citrate for 30 minutes at high temperature. Slides were cooled and washed with 2X saline sodium citrate (SSC). Tissues were pretreated with 3□g/ml proteinase K in 0.2 N hydrochloric acid for 10 minutes at 37 °C and washed with 2X SSC. Tissues were treated with lysozyme (2 mg/ml) for 30 minutes at 37 °C and incubated with 0.1% Triton-X-100 in PBS for 5 minutes at room temperature as done in ^71^. Following pretreatment, tissues were dehydrated in ethanol, air-dried, and tissue DNA was denatured in 50-70% formamide (Sigma, St. Louis, MO) in 2x SSC for 30 minutes at 65 °C. 50 ng/□l of the fluorescent probes in the hybridization buffer was deposited onto tissue at incubated in a moist HybEZ II Oven (Advanced Cell Diagnostics Newark, CA) from 3h to overnight at 45 °C. The following day, tissue was washed in 2X SSC and counterstained with ProLong Diamond Antifade Mountant with DAPI (Invitrogen, Eugene). Alternatively, tissues were formalin fixed, cryopreserved in 15% and 30% sucrose, and embedded in Tissue-Tek O.C.T Compound (Sakura Finetek, Torrance CA) prior to freezing. 5-micron sections were hydrated in sterile PBS, pretreated, and hybridized as described with FPPE tissues. To reduce tissue, auto-fluorescence for both immunofluorescence and FISH, sections were treated with Vector TrueVIEW (Vector Laboratories, Burlingame CA) and washed prior to sealing with the antifade.

### Immunofluorescence, transmission electron microscopy, and brightfield

Formalin-fixed paraffin embedded (FPPE) term BP biopsies and cultured explants challenged with *R. insidiosa* were processed in 3-5 micron sections and underwent H&E as described previously. Immunofluorescence staining was performed as described in ^41,72^ with a mouse monoclonal antibody to HLA-G (1:500, Santa Cruz Biotechnology, Santa Cruz CA) and a species-specific Alexa Fluor −594. Sections and cells were imaged using the Zeiss confocal microscope using 40x-63x oil immersion. H&E stains of BP biopsies, cultured explants, and mouse placentas were imaged using the brightfield setting of the pathology slide scanner, Nanozoomer (Hamamatsu Photonics), or the Eclipse E3800 Digital Microscope. For the Nanozoomer, images were analyzed using the NDP.view2 viewer software. For the Eclipse E3800 Digital Microscope, images were analyzed or using DPController (Olympus v. 3.2.1.276). Transmission electron microscopy for *R. insidiosa*-challenged JEG-3 cells and cultured explants was conducted as described in ^72^.

### JEG-3 adherence and invasion assay

JEG-3 cells were obtained from ATCC (HB-36) and cultured as described in ^73^. Cells were challenged with MJ602 at a multiplicity of infection of 0.01, 0.1, and 1. Plates were centrifuged at 1500 RPM for 5 minutes and incubated for 8h. At each MOI, wells were washed three times with warm sterile PBS. Percent adherence and invasion was determined as described in ^74^ with the following modifications: to quantify cellular adherence and invasion of bacteria, wells were scraped in 900 µL of PBS and placed in a centrifuge tube. 100 µL of 1% Triton-X-100 was added to cells, incubated, and then homogenized through gentle pipetting. Total bacteria per well was determined by adding sterile PBS to wells up to a volume of 900□L, which was followed by the addition of 100 µL of 1% Triton-X for 10 minutes. Cells in the well were homogenized through gentle pipetting. Homogenates were serially diluted and plated onto R2A and incubated at 32 °C overnight. CFUs for adhered and invaded bacteria and total bacteria were completed in triplicate for each MOI. Percent bacterial adherence and invasion was calculated as follows: CFUs of adhered / CFUs of total bacteria × 100 (n=4).

### Lactate dehydrogenase (LDH) assay

5×10^4^ JEG-3 cells were seeded onto 24-well plates. Cells were challenged with either *R. insidiosa* (MJ602) or *L. monocytogenes* (EGD) for 8 h at MOI of 0.01, 0.1, and 1 in triplicate. Cell lysis upon *R. insidiosa* challenge was measured using the CytoTox 96 Non-Radioactive Cytotoxicity kit (Promega, Madison, WI USA), which quantifies lactate dehydrogenase release in the culture media ^75^, according to the manufacturer’s instructions. Percent cytotoxicity was calculated according to the manufacturer’s instructions. Percent cytotoxicity at each MOI was plotted (n=3) using Graphpad Prism (v. 8).

### cDNA synthesis, RT-qPCR, and species-specific qPCR

Total mRNA was isolated with TRIzol (Thermo Fisher Life Technologies Corporation, Carlsbad CA) and treated with DNase I (Life Technologies Ambion, Carlsbad CA). RNA quality was validated with 260/280 ratio above 1.80. Only samples with a ratio over 1.80 were transcribed into cDNA. Complementary DNA (cDNA) was synthesized from 1 μg of total RNA by using SuperScript II Reverse Transcriptase (Thermo Fisher Scientific, Life Technologies, Carlsbad CA) according to the manufacturer’s instructions. 140 µL of water was added to the cDNA product. Each 20 µL qPCR reaction sample contained 10 µL of 2X SsoAdvanced Universal SYBR Green Supermix, 2 µL of cDNA, 7 µl of H_2_O, and 0.5 µL (10 µM) of each of forward and reverse primer. The NF-κB (5’-3’) primer pairs used in this study were published previously and include the forward primer AACAGAGAGGATTTCGTTTCCG (Primerbank ID: 259155300c1) and the reverse primer TTTGACCTGAGGGTAAGACTTC (PrimerBank ID: 259155300c1). The yfp primer pairs used in this study are presented in **Table S3.** *R. insidiosa*-specific PCR primer pairs (5’-3’) were published previously and include the forward primer Rp-F1 (ATGATCTAGCTTGCTAGATTGAT) ^76^ and the reverse primer R38R1 (CACACCTAATATTAGTAAGTGCG) ^33^. Primers were generated commercially (Integrated DNA Technologies, Coralville). The CFX96 Touch Real-Time PCR Detection System (Bio-Rad Laboratories) was used. qPCR consisted of 40 cycles as follows: 5 s at 95°C (30 s for the first cycle only), 30 s at 60°C, 31 s at 65°C; followed by 5 s at 65°C. The transcript level in each sample was measured in duplicate, and the melt curves were analyzed to confirm product specificity. The fold-change of unchallenged compared to *R. insidiosa*-challenged JEG-3 cells was determined at each MOI.

### Cytokine Secretion

Media collected post-*R. insidiosa* challenge (for 8 hours at an MOI 0, 0.01, 0.1, and 1) were profiled for pro-inflammatory cytokines including IL-6, IL-8, and TNF-α using the Milliplex Map Kit Human Cytokine/Chemokine Magnetic Bead Panel (EMD Millipore, Billerica). The procedure was followed according to the manufacturer’s instructions.

### Human Microbiome Project (HMP) dataset

The HMP Community Profile data page provides 16S sequencing datasets sampled from multiple body sites from healthy individuals. We opted to use the preprocessed HMPv35 dataset (HMPv35.R.data), which includes variable regions 3 through 5 in the binary file format, using http://joey711.github.io/phyloseq-demo/HMP_import_example.html instead of importing the dataset directly from the public HMP data files. R (v. 3.5.2) was used to analyze the HMP dataset and is available upon request. Briefly, a subset dataset was generated comprising non-contaminated, correctly labeled, female-derived samples. The *Ralstonia* genera was subset from this dataset. The following packages were used: Phyloseq (v. 1.24.2), stringi (v. 1.4.3). The abundance of Ralstonia-specific OTUs in the saliva, throat, hard palate, and stool were plotted using Graphpad Prism (v. 8).

### Mouse experiments

Vaginal smears were monitored in 8 to 15 week-old nulliparous female wild-type mice obtained from Jackson Laboratory (Bar Harbor, Maine). At estrus stage of their cycles, females were mated with WT males at a female-male ratio 1:1, and day 0.5 of pregnancy was defined as the first observation of a vaginal plug. All mice were on a C57BL6 background and housed under a 12-h light/12-h dark cycle in mouse pathogen-permitted breeding facility. Mice were challenged at day 15.5 via intrauterine injection and oral gavage with *R. insidiosa* yfp-strain GGW102. For oral gavage, 100 µL harboring 1×10^6^ (n=3) or 1×10^8^ (n=3) CFUs of *R. insidiosa* (GGW102) was administered using a 1 mL syringe and gavage needle. SHAM mice were used as controls (n=4). For intrauterine injections, dams were anesthetized using isoflurane and oxygen. Under sterile conditions, the abdomen was prepped with betadine and saline and a vertical skin incision was made while avoiding mammary glands. For intrauterine injection, 100 µL harboring 1×10^3^ (n=5) CFUs of were administered to the uterus with a 1 mL syringe at two sites between 1^st^ and 2^nd^ fetus. 50 µL was administered at each injection site, without entering the amniotic cavity. Dams were allowed to recover following routine closure with absorbable Vicryl for peritoneal cavity ligation. SHAM-operated mice were used as controls for intrauterine injected mice (n=3). Pregnant dams were monitored for signs of labor and sacrificed at day 18.5 if there were no signs of labor. Upon sacrifice, pregnancy tissues were collected including uteri and myometria, placentas, and fetal membranes. Feto-placental units were underwent formalin-fixation and paraffin embedding or were formalin-fixed and frozen as described previously. Additionally, placental weight, fetal length, fetal width, and fetal weight were determined and fetal membranes, placentas, and uteri/myometria were quantified for CFU/g.

### Recovery and detection of *R. insidiosa* in the tissues of challenged mice

Placentas, fetal membranes, and uteri/myometria were collected from feto-placental units and were washed PBS and weighed. Tissues were homogenized in 100 - 500 µL of R2B using sterile beads for 8 minutes at high speed. The homogenate was serially diluted and plated on R2A in triplicate. CFU/g tissue was calculated and log transformed. If round white colonies were cultivated from the fetal membrane, placentas or uteri/myometria, the presence of *R. insidiosa* was confirmed using species-specific FISH on at least two feto-placental units from 2 dams. Briefly, one colony was smeared and heat-fixed onto a glass slide and hybridized with the *R. insidiosa*-specific probe and DAPI as described previously. H&E was performed on FPPE tissues and FISH targeting *R. insidiosa* was performed on fixed-frozen placentas.

### Statistics

For mouse studies, the mean ± SEM measurements from multiple feto-placental units were calculated and the data was presented in graphs as the median measurement per dam. The Mann-Whitney *U* test was performed for comparisons between two groups, and the Kruskal-Wallis test was performed with Dunnett’s post-test for multiple comparisons. Graphpad Prism 8.0 was used for all analyses. *P* < 0.05 was considered significant.

## Acknowledgments

We thank the Center for the Women and Infants’ Health Specimen Consortium at Washington University for sample collection; Wandy Beatty and the Molecular Microbiology Imaging Facility for assistance with TEM; and Deborah Frank for comments on the manuscript. This work was supported by a Preventing Prematurity Initiative grant from the Burroughs Wellcome Fund, a Prematurity Research Initiative Investigator award from the March of Dimes (to I.U.M), the NASA cooperative agreement NNX16ZHA001C (M.J.W), the NIH grant 2T32GM007067-42 (L.A.P), the NIH Grant T32 HL076122 (G.W.W), and the NIH CTSA Grant (UL1 TR0004488). The *L. monocytogenes* EGD strain was provided as a gift from Dr. Brian Edelson.

The research reported in this project was also supported by the Washington University Institute of Clinical and Translational Sciences grant UL1TR002345 from the National Center for Advancing Translational Sciences (NCATS) of the National Institutes of Health. The content does not necessarily represent the official view of the NIH.

## Additional Information

### Contributions

L.A.P and I.U.M. designed the research; M.J.W. provided the MJ602 (WT) strain, generated YFP-labeled *R. insidiosa*, and generated the *R. insidiosa-*specific fluorescent in situ hybridization probes. L.A.P, G.G.W, C.S.J, and Y.Y. conducted experiments. L.A.P and I.U.M. analyzed data and wrote the manuscript. All authors have read and approved the manuscript.

