## Supplementary Materials for "Functional characterization of *Ralstonia insidiosa*, a bona fide resident at the maternal-fetal interface"

### Supplemental figures and tables

#### Supplementary tables

Table S1. **Overview of the 16S rRNA fluorescent in situ hybridization probes applied to basal plate specimens in this study.**

Table S2. **Overview of the isolates used.**

Table S3. **Primer pairs targeting yellow-fluorescent protein.**

#### Supplementary figures

Figure S1. **Evaluation of probe functionality by whole bacterial cell fluorescent in situ hybridization using a *R. insidiosa* WT strain (MJ602).**

Figure S2. **Evaluation of FISH probe functionality and yfp-expression using GGW102.**

Figure S3. **Growth profiles *R. insidiosa* strains MJ602 and GGW102 in monocultures.**

Figure S4. **Placentas and fetuses from mice challenged with *R. insidiosa* do not exhibit significant anatomical anomalies.**

| Probe name | Sequence 5’ → 3’ | Label (5’ end) | Specificity | Source |
|---|---|---|---|---|
| EUB338 | GCTGCCTCCCGTAGGAGT | CY3 | Conserved domain of the eubacterial 16S rRNA gene | Amann et. al 1990 [67] |
| NON-EUB338 | ACTCCTACGGGAGGCAGC | CY3 | Nonsense control complementary to EUB338 | Wallner et. al 1993 [68] |
| *R. insidiosa* | TTAGTAAGTGCGATTTCTTTCCGGA | FITC | Species-specific sequence of *R. insidiosa* 16S rRNA gene | This study |

Table S1. **Overview of the 16S rRNA fluorescent in situ hybridization probes applied to basal plate specimens in this study.**

| ***R. insidiosa* isolate** | **Source** | **Growth conditions** |
|---|---|---|
| MJ602 (WT) | Johnson Space Center as *R. insidiosa* 130770013-1 [73] | Static; 32 °C Reasoner's 2B Medium |
| GGW102 (YFP strain) | attTn7 integration of *yfp* into strain MJ602 | Static; 32 °C Reasoner's 2B Medium |

Table S2. **Overview of the isolates used.**

| Primer Name | Forward (5’-3’) | Reverse Forward (5’-3’) | Product Length (bp) |
|---|---|---|---|
| YFP 1 | GCTACCCCGACCACATGAAG | AAGAAGATGGTGCGCTCCTG | 82 |
| YFP 2 | ACGTAAACGGCCACAAGTTC | CTTCATGTGGTCGGGGTAGC | 176 |
| YFP 3 | AGCAAAGACCCCAACGAGAA | TCGTCCATGCCGAGAGTGAT | 83 |
| YFP 4 | GAACCGCATCGAGCTGAA | TGCTTGTCGGCCATGATATAG | 111 |

Table S3. **Primer pairs targeting yellow-fluorescent protein.**

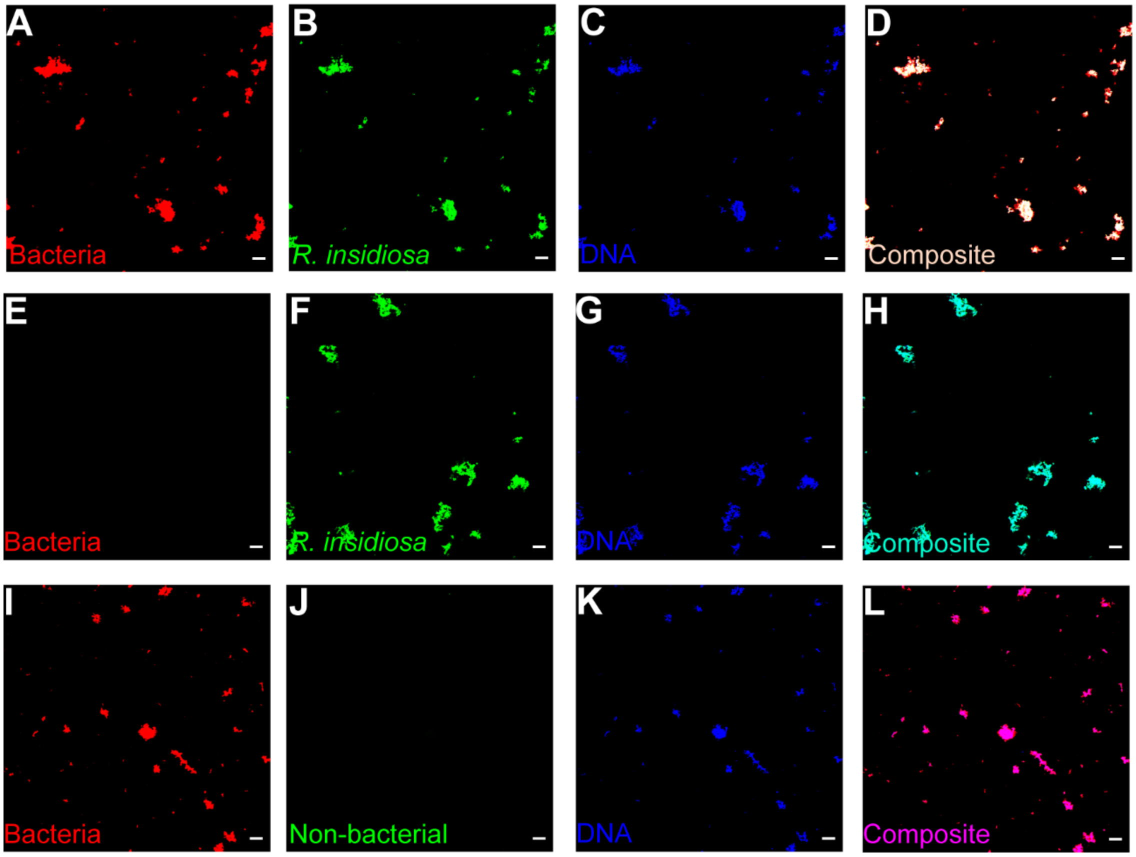


Figure S1. **Evaluation of probe functionality by whole bacterial cell fluorescent in situ hybridization using a *R. insidiosa* WT strain (MJ602).** EUB338-CY3 was used as a universal bacterial probe (bacteria-red), *R. insidiosa*-FITC was used as a species-specific probe (*R. insidiosa*-green), NON-EU338-FITC represents the complement to the universal eubacterial probe and was used as a negative control probe (non-bacteria-green), and DAPI was used for host nuclear and bacterial DNA staining (DNA-blue). Whole bacterial cells were simultaneously hybridized with (A-D) bacterial probe, *R. insidiosa* probe (*R. insidiosa*-green) and DAPI (DNA-blue); (E-H) *R. insidiosa* and DAPI; or with the bacterial and non-bacterial probe. Fluorescent images were acquired separately using thee filter sets (CY3, FITC, DAPI) are displayed as a final image (composite) (Magnification 40x oil, scale bar:  20 µm).

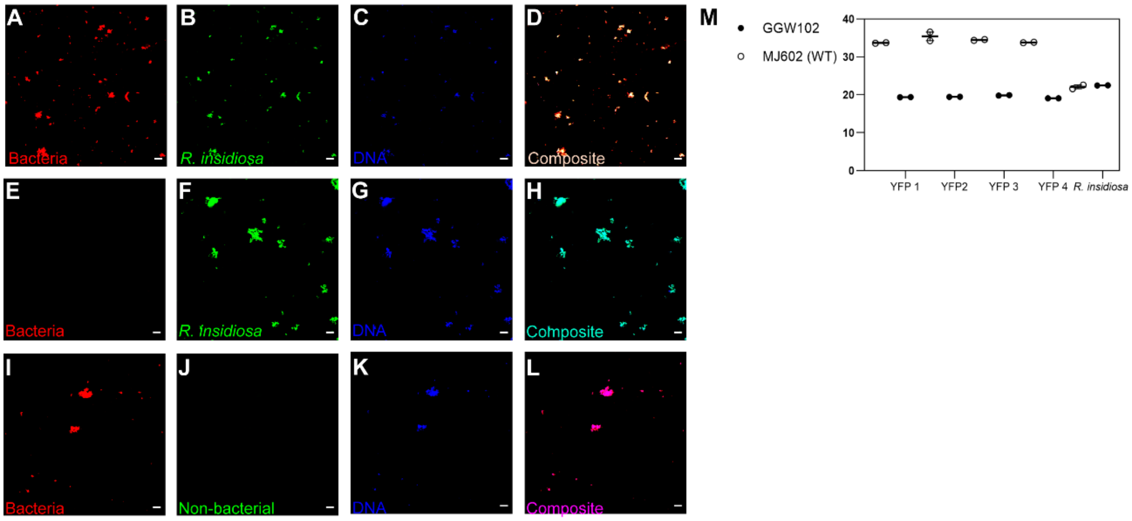


Figure S2. **Evaluation of FISH probe functionality and yfp-expression using GGW102.** Whole bacterial cells were simultaneously hybridized with the universal bacterial probe (bacteria-red), *R. insidiosa* probe (*R. insidiosa*-green) and DAPI (DNA-blue); (E-H) the *R. insidiosa* probe and DAPI; or with the bacterial probe (red) and non-bacterial probe (green). Fluorescent images were acquired separately using thee filter sets (CY3, FITC, DAPI) are displayed as a final image (composite) (Magnification 40x oil, scale bar: 20 µm). (M) YFP primers (YFP 1-4) were used verify the presence of YFP. YFP is expressed in GGW102, but not in MJ602. The *R. insidiosa*-specific product is amplified in both isolates. Cq values are reported.

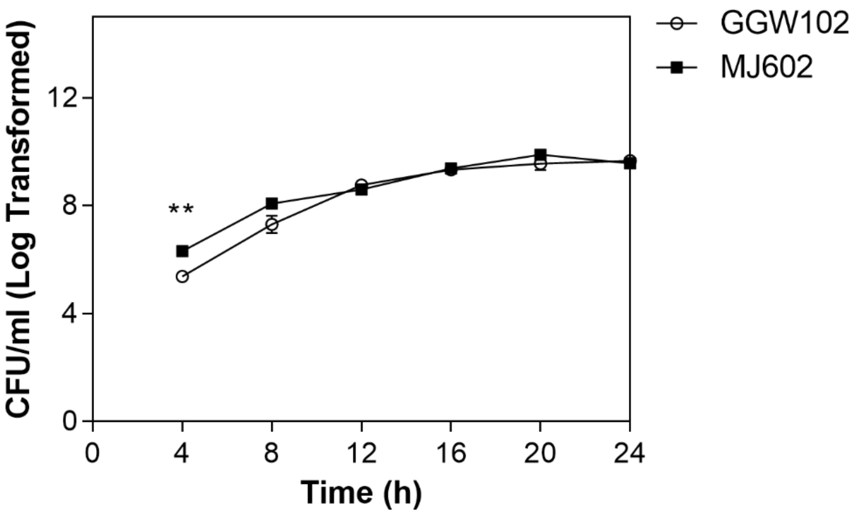


Figure S3. **Growth profiles *R. insidiosa* strains MJ602 and GGW102 in monocultures.** The data is the average CFUs over the course of 24 hours for MJ602 and GGW102 (n = 6 colonies each). Single colonies were inoculated in 20 mLs of R2B. Although there is a significant difference in the log CFUs in MJ602 compared to GGW102 at 4 h ($p = 0.00220$), there as no significant difference between the growth of isolates at 8h and onward (8h, p =0.180, 12h, p = > 0.999; 16h, p = >0.999, 20h, p = 0.240, 24h, p = 0.818) suggesting that they can be used interchangeably for replication analysis ex vivo, in vitro, and in vivo studies. P-values were calculated using the two-tailed Mann-Whitney U test at each time point. ** indicates a $p < 0.005$.

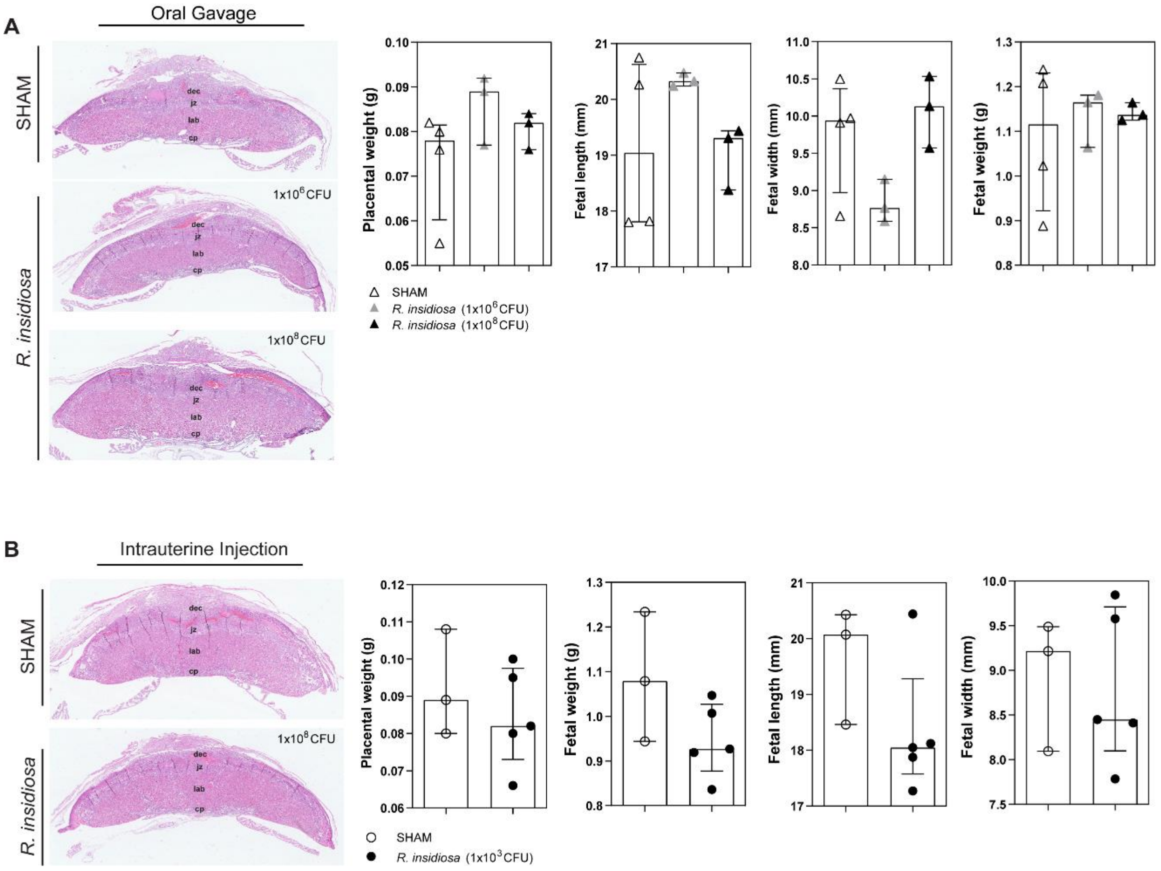


Figure S4. **Placentas and fetuses from mice challenged with *R. insidiosa* do not exhibit significant anatomical anomalies.** (A) Representative hematoxylin and eosin staining of mouse placentas (gd 18.5) derived from SHAM (n= 4) or *R. insidiosa* challenged dams following oral gavage of 1x10$^6$ (n=3) and 1x10$^8$ (n=3) CFUs at gd 15.5. Fetoplacental units of *R. insidiosa*-challenged dams compared to SHAM controls exhibit no differences in the median placental weight (n=10, Kruskal-Wallis statistic = 2.92; p = 0.260), fetal length by oral gavage (n=10, Kruskal-Wallis statistic = 2.38; p = 0.326), fetal width (n=10, Kruskal-Wallis statistic = 4.27; p = 0.123), or fetal weight (n=10, Kruskal-Wallis statistic = 0.164; p = 0.941). (B) Representative hematoxylin and eosin staining mouse placentas (gd 18.5) from (H) SHAM or (I) *R. insidiosa*-challenged dams following intrauterine injection of 1x10$^3$ CFUs at gd 15.5. Fetoplacental units of *R. insidiosa*-challenged dams compared to SHAM controls exhibit no differences in the median placental weight (Mann-Whitney U = 5.50; p = 0.643), fetal length by oral gavage (Mann-Whitney U=3; p =0.250), fetal width (Mann-Whitney U=7; p = >0.999), or fetal weight (Mann-Whitney U = 2; p = 0.143). P-values were calculated using the two-tailed Mann-Whitney U test.
